# Moderate prenatal alcohol exposure alters GABAergic transmission and the actions of acute alcohol in the CeM of adolescent rats

**DOI:** 10.1101/2024.08.16.608070

**Authors:** Sarah E. Winchester, Marvin R. Diaz

## Abstract

Individuals with prenatal alcohol exposure (PAE) are at a higher risk for developing alcohol use disorder (AUD). Using a rat model of PAE on gestational day 12 (G12; ~2^nd^ trimesters in humans), a critical period for amygdala development, we have shown disruptions in medial central amygdala (CeM) function, an important brain region associated with the development of AUD. In addition to this, acute ethanol (EtOH) increases GABA transmission in the CeM of rodents in a sex-dependent manner, a mechanism that potentially contributes to alcohol misuse. How PAE alters acute alcohol’s effects within the CeM is unknown. Given these findings, we investigated how PAE may interact with acute alcohol to alter neuronal and synaptic mechanisms in the CeM of adolescent rats in order to understand PAE-induced alcohol-related behaviors. Under basal conditions, PAE males showed reduced rheobase, indicative of reduced excitability, and females showed a reduction in GABA transmission, indicated by lower spontaneous inhibitory postsynaptic currents (sIPSCs). We found that acute EtOH increased sIPSCs in control males at a moderate concentration (66 mM), while PAE males showed increased sIPSCs only at a high concentration (88 mM). Adolescent females, regardless of PAE status, were largely insensitive to EtOH’s effects at all tested concentrations. However, PAE females showed a significant increase in sIPSCs at the highest concentration (88 mM). Overall, these findings support the hypothesis that PAE leads to sex-specific changes in synaptic activity and neuronal function. Future research is needed to better understand the specific mechanisms by which acute EtOH’s affects neurotransmission in the adolescent brain of individuals with a history of PAE.

## 1.0 Introduction

It is well known that prenatal alcohol exposure (PAE) can lead to severe adverse consequences in offspring, termed fetal alcohol spectrum disorder (FASD). The CDC estimates that for every 1,000 births, 0.2 to 7 infants are born with FASD in the U.S. (Dejong et al., 2019). Individuals with FASD can present with a range of physical, cognitive, and emotional impairments, including future complications with the misuse of alcohol, beginning in adolescence and persisting throughout the lifespan (Baer et al., 1998; Baer et al., 2003). Importantly, offspring exposed to alcohol early in gestation are 2.95 times more likely to develop alcohol use disorder (AUD) by the time they reach 21 years of age (Alati et al., 2006).

Our lab has characterized a gestational day 12 (G12; equivalent to ~2nd trimester in humans) moderate PAE (mPAE) rat model. G12 is a crucial time period for the development of the amygdala as amygdalar neurons form from the neuroepithelium and migrate to assemble the amygdalar subdivisions between G10 and G13 (Soma et al., 2009). More specifically, formation of the medial nucleus of the central amygdala (CeM) occurs, a brain structure implicated as a key brain region involved in the misuse of alcohol along with the development of an alcohol use disorder (Gilpin et al., 2015; Hyytia and Koob, 1995; Roberto et al., 2003). Using our mPAE model, we have observed disruption in the function of the CeM; more specifically, we found an increase in GABA transmission onto and decrease in excitability of CeM neurons of mid-adolescent male rats (Rouzer and Diaz, 2022). However, how PAE may directly alter neurotransmission in this region as a mechanism contributing to alcohol-related behaviors is not well understood.

Acute application of ethanol (EtOH) to CeM neurons increases GABA transmission in adult naïve male rats, with a majority of the cells responding to a 44 mM concentration of EtOH (Kirson et al., 2021; Roberto et al., 2003; Roberto et al., 2004). This increase in GABA transmission in response to EtOH occurs through a presynaptic mechanism, with an increase in GABA release, in addition to some adaptations in postsynaptic function (Roberto et al., 2003). Interestingly, a dose response curve (11-88 mM) revealed that adult naïve female rats are largely insensitive to the actions of acute EtOH on GABA transmission, whereas adult naïve males showed the largest increase at moderate to high concentrations (44-88 mM) (Kirson et al., 2021). Furthermore, an increase in GABA transmission was only observed in alcohol-dependent adult females at the highest tested concentration (88 mM), and in alcohol-dependent males at moderate concentrations (44-66 mM) (Kirson et al., 2021). These findings suggest that GABA transmission within the CeM in adult females is largely insensitive to the actions of acute EtOH, and that chronic alcohol exposure produces sex-dependent alterations in CeA responsivity to acute EtOH in adulthood. These sex differences may contribute to differences observed in alcohol intake not only in rodents, but in adolescent humans as well (Becker and Koob, 2016; CDC, 2024).

Importantly, individuals with PAE have a propensity to misuse alcohol starting in adolescence, similar to that seen in alcohol-dependent individuals in adulthood. However, to our knowledge, there has been no research directly examining the effect of acute EtOH on GABA transmission within the CeM specifically in adolescence nor how PAE alters the actions of acute EtOH. To understand the mechanisms contributing to PAE-induced behaviors, it is necessary to investigate the impact of acute EtOH on neuronal and synaptic function in structures associated with the observed behaviors. Therefore, this study aimed to identify how PAE alters the actions of EtOH on GABA transmission in the CeM of adolescent male and female offspring. We hypothesized that G12 mPAE would produce sex-dependent alterations in CeM neuronal function, resulting in an augmented increase in GABA transmission induced by acute EtOH in adolescent males. Given that GABA transmission is increased in alcohol-dependent adult females at highert concentrations, we hypothesized that we would see a similar effect in PAE adolescent females.

## 2.0 Materials and Methods

### 2.1 Animals

Male and female Sprague Dawley breeders were purchased from Envigo (Indianapolis, IN, USA) for in-house breeding. Animals were pair housed and acclimated to the colony room for two weeks after arrival. Animals had *ad libitum* access to standard chow (Purina 5LOD rat chow laboratory diet) and water and were maintained in a temperature-controlled colony (22°C) on a 12:12h light/dark cycle (lights on from 0700 to 1900 hours). All electrophysiology experiments were performed in late-adolescent offspring (postnatal day (P) 45-P55) from litters produced in our colony. All experiments were approved by Binghamton University’s Institutional Animal Care and Use Committee (IACUC).

### 2.2 Breeding

Two females and one male were housed together over the course of four days. Each morning, females were checked for pregnancies via vaginal smears as previously described (Mooney and Varlinskaya, 2011; Rouzer et al., 2017; Rouzer and Diaz, 2022). When sperm was detected (G1), females were single housed with *ad libitum* access to breeder chow (Purina 5008 rat chow laboratory diet) and water. Dams were weighed throughout gestation (G1, G10, G20) to monitor viability of pregnancy. After parturition, all pups were left with the mother for 2 days before culling began. All litters were culled to 12 pups - when possible, a 1:1 male to female ratio was maintained. At day 21, pups were weaned and pair housed with same-sex litter mates until experimental testing began.

### 2.3 Moderate Prenatal Alcohol Exposure (mPAE)

Pregnant Sprague Dawley rats were put in fresh cages and transferred to vapor chambers on G12. Animals were exposed to either vaporized ethanol (95% ethanol) or room air for 6 hours as previously described (Przybysz et al., 2023; Rouzer et al., 2017; Rouzer and Diaz, 2022). EtOH vapor concentration levels were checked throughout the exposure by extracting air from the EtOH chamber through an access port (1:30 dilution with room air) and read with a breathalyzer (Intoximeters), targeting vapor concentration levels of 5.7 – 6.5 g/L. Animals had access to food and water during exposures. After 6 hours, animals were transferred to a clean cage, and food and water was replaced to prevent further EtOH exposure. Animals were left undisturbed until parturition.

### 2.4 Blood Ethanol Concentration (BECs)

To ensure that EtOH vapor chambers and breathalyzer were working properly, a different subset of pregnant dams was exposed to vaporized EtOH and tail blood collection (via heparinized Natelson blood collecting tubes) was performed every 2 hours until the end of the 6-hour exposure. All blood samples were centrifuged for 15 minutes and plasma was extracted for BEC testing. BECs were measured using an Analox AM1 Alcohol Analyser against a 50 mg/dL ethanol standard (Analox Instruments Ltd, The Vale, London).

### 2.5 Slice Electrophysiology

#### 2.5.1 Slice Preparation

Slices were obtained with procedures previously described by our lab (Rouzer and Diaz, 2021, 2022). Rats were anesthetized with 5% isoflurane, quickly decapitated and brains were rapidly removed and placed in oxygenated (95% O2, 5% CO2) sucrose artificial cerebrospinal fluid (sucrose ACSF) containing: 2 mM KCl, 1.3 mM NaH_2_PO_4_, 26 mM NaHCO_3_, 220 mM sucrose, 10 mM glucose, 12 mM MgSO_4_, 0.2 mM CaCl_2_, and 0.42 mM ketamine. Using a Vibratome (Leica Microsystems. Bannocknurn, IL, USA), coronal slices (300 μm thick) containing the CeM were obtained and incubated for at least 40 minutes in 33°C oxygenated artificial cerebrospinal fluid (ACSF) containing: 125 mM NaCl, 2 mM KCl, 1.3 mM NaH_2_PO_4_, 26 mM NAHCO_3_, 10 mM glucose, 2 mM CaCl_2_, 1 mM MgSO_4_, 0.4 mM ascorbic acid. All experiments were performed ~1-5 h after slice preparation.

#### 2.5.2 Whole-cell patch clamp electrophysiology

Brain slices containing the CeM were transferred to the recording chamber and submerged in oxygenated ACSF at 32°C which was superfused over the slice at 3.3 ml/min as previously described (Rouzer and Diaz, 2021, 2022). The CeM was identified using the Allen Brain Atlas as a guide, and neurons within the CeM were visually identified. Only neurons with a membrane capacitance of ~50 pF were used for recordings. Recording electrodes with a tip resistance of 3-5 MΩ were pulled from borosilicate glass capillary tubing (Sutter Instruments) using a Flaming-Brown puller (Sutter Instruments). Recording electrodes were filled with KCl (7.25 pH with KOH) containing: 135 mM KCl, 10 HEPES, 2 mM MgCl_2_, 0.5 mM EGTA, 5 mM Mg-ATP, and 1 mM Na-GTP; 300 mOsm. Cells were patched and opened in voltage clamp. Before recordings began, neurons equilibrated for 5 minutes. A MultiClamp 700B (Molecular Devices, Sunnyvale, CA) at 10 kHz, filtered at 1 kHz was used for electrophysiology data collection, then stored using pClamp Software (Molecular Devices) for analysis. Spontaneous inhibitory postsynaptic currents (sIPSC) were analyzed using MiniAnalysis (Synaptosoft Inc.).

#### 2.5.3 Current Clamp

Before current-clamp experiments were conducted, the cell was switched into a current neutral mode to assess resting membrane potential (RMP). Current-clamp mode was used to assess measures of neuronal excitability. Previous work in our lab identified a PAE-induced depolarization of the RMP in CeM neurons (Rouzer and Diaz, 2022), therefore, for current step injections, cells were forced to a membrane potential of −70mV. Recordings were stored for analysis of rheobase (time to first action potential) and # of action potentials (APs) induced by current steps (increments of 10 pA, 500ms duration). The first AP elicited in response to the lowest current injection was analyzed for AP threshold, amplitude, and half-width.

#### 2.5.4 Voltage clamp

For voltage clamp recordings, to isolate and record GABA_A_-mediated sIPSCs, AMPA and NMDA receptors were pharmacologically blocked using 1 mM kynurenic acid and 50 μM APV, respectively. To assess the effects of EtOH on sIPSC frequency and amplitude from CeM neurons, EtOH (44, 66, or 88 mM) was applied to separate brain slices for at least 10 minutes. Only recordings with access resistance change <20% were used for analysis.

### 2.6 Statistics

All statistical analyses were performed using GraphPad 9 software (Prism). No more than 1 cell was recorded per animal for any experiment. Based on previously reported sex-differences in PAE and acute ethanol effects on CeM neurophysiology (Kirson et al., 2021; Rouzer and Diaz, 2021, 2022), all electrophysiological data was separated by sex *a priori* for statistical analyses. All data were analyzed for normal distribution using Kolmogorov-Smirnov test and outliers were statistically identified using Grubb’s test in Prism. Data that were not normally distributed were analyzed using non-parametric tests. RMP, rheobase, baseline sIPSC frequency and amplitude was analyzed using an unpaired t-test between Air and PAE groups. A within cell analysis was conducted using a one-sample t-test to examine drug effects on sIPSC frequency and amplitude relative to baseline within each group. A 2-way ANOVA was used to analyze firing experiments and dose response curves. Sidak’s multiple comparison tests were used to determine specific group differences in the event of significance. Significance was defined as p ≤ 0.05. All data is presented with the standard error of the mean.

## 3.0 Results

### 3.1 Dam Blood Ethanol Concentrations (BECs)

BECs were collected and analyzed every 2 hours over the 6-hour exposure period, with the exposure starting at hour 0. The peak BEC was 108.3 ± 52.20 (n=4; **Table 1**), occurring at the last (6th) hour of exposure. This exposure is considered a mPAE, as previously determined (Valenzuela, 2012).

**Table 1.** Average blood ethanol concentration (BEC) time course. A subset of dams (n=4) were used to determine BECs over the course of the 6-hour exposure.

| Hour | Mean (mg/dL) | Std. Error of Mean |
| --- | --- | --- |
| 0 | 5.70 | 2.209 |
| 2 | 57.41 | 16.14 |
| 4 | 79.97 | 10.75 |
| 6 | 108.3 | 26.10 |

### 3.2 Membrane Properties

To assess how PAE may alter basal membrane properties, membrane resistance (Rm) and membrane capacitance (Cm) were monitored throughout entire recordings. No differences were observed in Rm or Cm for males (Rm: t (54) = 0.270, *p* = 0.788; Cm: t (54) = .0708, *p* = 0.482; **Table 2**) or females (Rm: t (56) = 0.527, *p* = 0.600; Cm: U = 396, *p* = 0.734; **Table 2**).

**Table 2.**
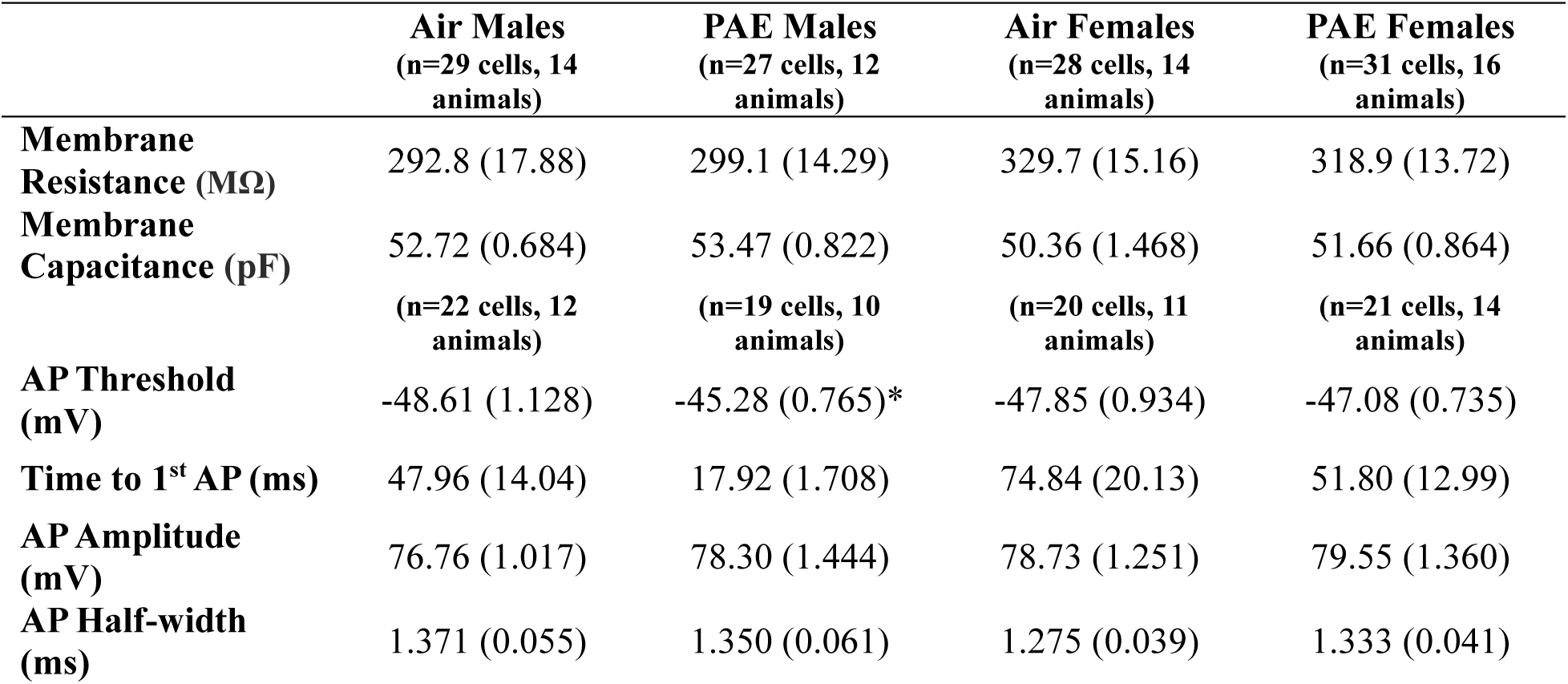
Membrane and action potential (AP) properties of CeM neurons, reported as mean (SEM). No differences were observed in membrane resistance (Rm), membrane capacitance (Cm), time to first AP, AP amplitude or AP half-width. However, PAE influenced AP threshold in males, where PAE male’s threshold was higher than air males. Females did not differ in this AP parameter. A test of normality was conducted before using the proper parametric test. * denotes statistical significance compared to control counterpart. * denotes p ≤ 0.05.

### 3.3 G12 mPAE influences the excitability of male and female CeM neurons

Resting membrane potential was assessed to investigate how mPAE may alter membrane properties. An unpaired t-test revealed no differences in resting membrane potential in males (t (45) = 1.674, *p* = 0.1012; **Figure 1A**) or females (t (44) = 0.241, *p* = 0.811; **Figure 1A**).

**Figure 1.**
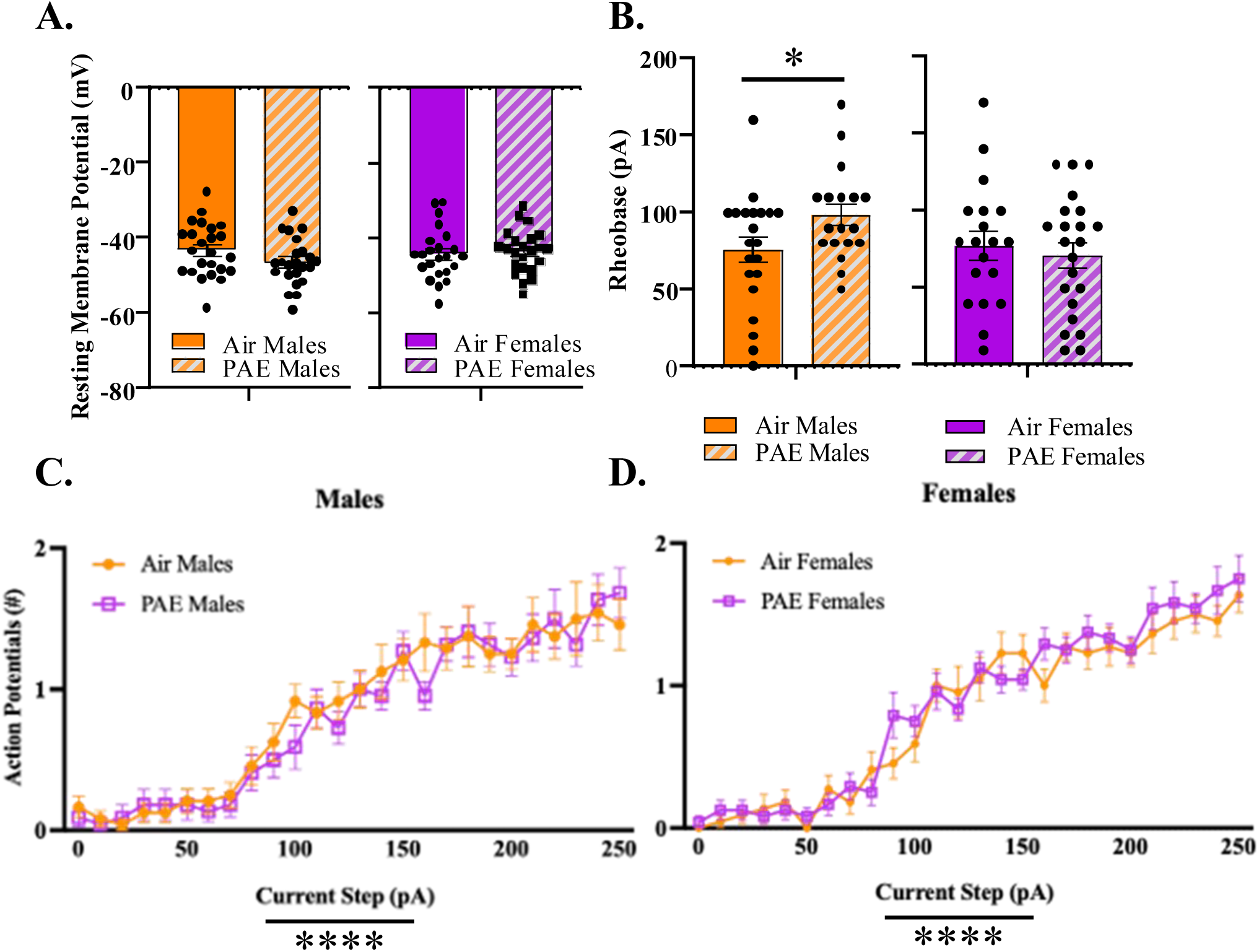
Membrane Properties and excitability of CeM Neurons. (**A**) We observed no differences in resting membrane potential (RMP) in males (air: n = 24 from 12 animals; PAE: n = 23 cells from 10 animals) or females (air: n = 23 cells from 13 animals; PAE: n = 23 cells from 15 animals). (**B**) There was a significant difference in the amount of current needed to elicit the first action potential in males (air: n = 21 cells from 12 animals; PAE: n = 19 cells from 10 animals). No differences were observed in rheobase for females (air: n = 18 cells from 13 animals; PAE: n = 22 cells from 15 animals). (**C**) We observed an effect of current step and no effect of exposure on the excitability of CeM neurons in males (air: n = 21 cells from 12 animals; PAE: n = 19 cells from 10 animals). (**D**) An effect of current step and exposure was observed in the excitability of CeM neurons in females (air: n = 18 cells from 13 animals; PAE: n = 22 cells from 15 animals). Test for normality were conducted before t-tests. A two-way ANOVA was used to test for differences in excitability. * denotes p ≤ 0.05. Bars represent standard error of mean.

We next examined how mPAE may alter excitability of CeM neurons. A portion of cells exhibited spontaneous AP firing making it impossible to assess current-induced excitability, therefore these cells were removed from the excitability analysis, but included in sIPSC analysis (air males: 4 cells, PAE males: 5 cells, air females: 9 cells, PAE females: 7 cells). For neurons that responded to current injection, the 1^st^ AP that occurred was assessed for the time it took to fire that AP, AP threshold, AP amplitude and AP half-width (**Table 2**). We observed a significant difference in the AP threshold for males (U = 110, *p* = 0.015), but not females (t (38) = 0.644, *p* = 0.524). In males and females, PAE did not affect the time to 1^st^ AP (males: U = 153, *p* = 0.229; females: U = 198, *p* = 0.767), AP amplitude (males: t (38) = 0.883, *p* = 0.383; females: t (39) = 0.443, *p* = 0.661), or AP half-width (males: U = 189, *p* = 0.789; females: t (39) = 1.020, *p* =0.314). When assessing rheobase, we found that mPAE males required more current to induce an AP (t (37) = 2.025, *p* = 0.050; **Figure 1B**), whereas no differences were observed in female rheobase (t (39) = 0.494, *p* = 0.624; **Figure 1B**). Current step injections were used to assess firing capabilities of cells. There was an effect of current steps observed in males (*F* (25, 1144) = 30.85, *p* < 0.0001) and females (*F* (25, 1144) = 54.60, *p* < 0.0001; **Figure 1C & 1D**), where the more current injected, the more AP were elicited. However, there was no effect of exposure on the excitability of cells in males (*F* (1, 624) = 0.618, *p* = 0.432; **Figure 1C**) or females (*F* (1, 1144) = 2.274, *p* = 0.132; **Figure 1D**).

### 3.4 G12 mPAE effects on basal GABA transmission

We examined how mPAE may alter spontaneous GABAergic transmission under basal conditions (**Figure 2**). We found no effect of PAE on sIPSC frequency in males (U= 391, *p* > 0.999; **Figure 2C**; depicted in exemplar traces **Figure 2A**). However, there was a significant decrease in basal sIPSC frequency in females (U = 278, *p* = 0.0412; **Figure 2C**; depicted in exemplar traces **Figure 2B**). We found no differences in the amplitude of sIPSCs in males (U = 354.5, *p* = 0.698; **Figure 2D**) or females (U =368, *p* = 0.424; **Figure 2D**) as a function of PAE.

**Figure 2.**
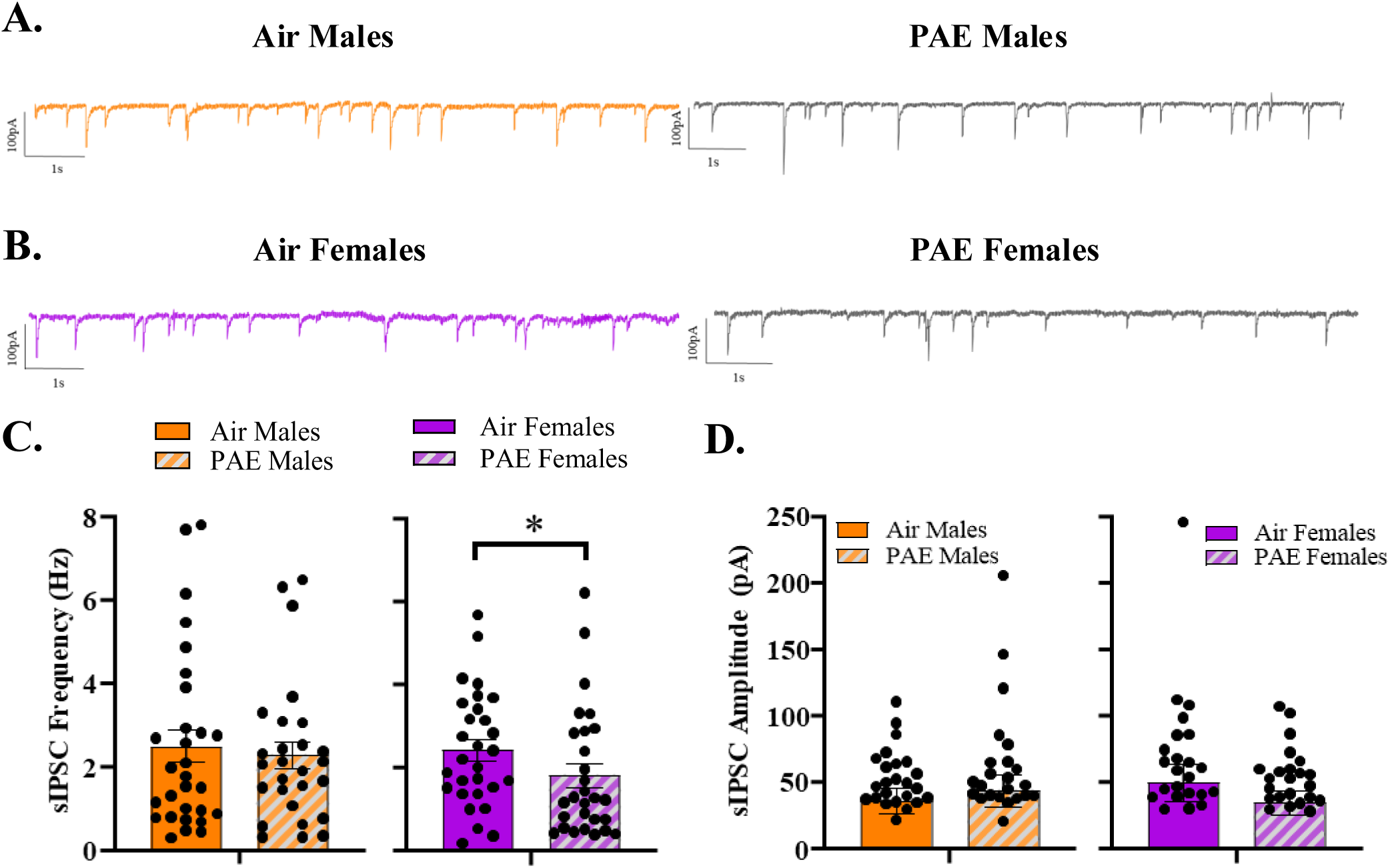
Basal spontaneous inhibitory post synaptic currents (sIPSCs). (**A**) Representative sIPSC activity of male CeM neurons: there was no difference in sIPSC frequency (**C**) or amplitude (**D**) (air: n = 29 cells from 14 animals; PAE: n = 27 cells from 12 animals). (**B**) Representative sIPSC activity of female CeM neurons: there was a significant effect of exposure on sIPSC frequency in females (**C**), with no difference in sIPSC amplitude (**D**) (air: n = 30 cells from 15 animals; PAE: n = 29 cells from 17 animals). All data was tested for normal distribution and the appropriate parametric tests were used. * denotes p ≤ 0.05. Bars represent standard error of mean.

### 3.5 G12 mPAE alters acute alcohol modulation of GABA transmission in males

To investigate how mPAE may alter acute EtOH actions on GABA transmission in the CeM of males, we first assessed the effects of EtOH (44, 66, & 88 mM) application on sIPSC frequency (**Figure 3**) - each concentration was applied to a single cell on separate slices. When examining changes in raw sIPSC frequency, we observed no changes at 44 mM EtOH in air (t (8) = 0.823, *p* = 0.434; **Figure 3A**) or PAE males (t (9) = 0.146, *p* = 0.887; **Figure 3A**). In air males, 66 mM EtOH significantly increased sIPSC frequency (t (9) = 3.000, *p* = 0.015; **Figure 3A**), however, this concentration of EtOH had no effect in PAE males (W = 25.00, *p* = 0.164; **Figure 3A**). Unexpectedly, there were no changes in sIPSCs following application of 88 mM EtOH in either air (t (9) = 1.271, *p* = 0.236; **Figure 3A**) or PAE males (t (7) = 1.924, *p* = 0.096; **Figure 3A**). Next, we analyzed drug effects by % change in activity from baseline. Consistent with observations in raw frequency, sIPSC frequency was significantly increased from baseline in air males at 66 mM (t (9) = 3.030, *p* = 0.014; **Figure 3B**), but not at 44 mM (t (8) = 1.009, *p* = 0.343; **Figure 3B**) or 88 mM (t (9) = 1.816, *p* = 0.103; **Figure 3B**). PAE males did not exhibit changes in sIPSC frequency at 44 mM (t (9) = 0.569, *p* = 0.583; **Figure 3B**) or 66 mM (t (8) = 1.715, *p* = 0.125; **Figure 3B**); however, we detected a significant increase in sIPSC frequency from baseline at 88 mM (t (7) = 2.685, *p* = 0.031; **Figure 3B**).

**Figure 3.**
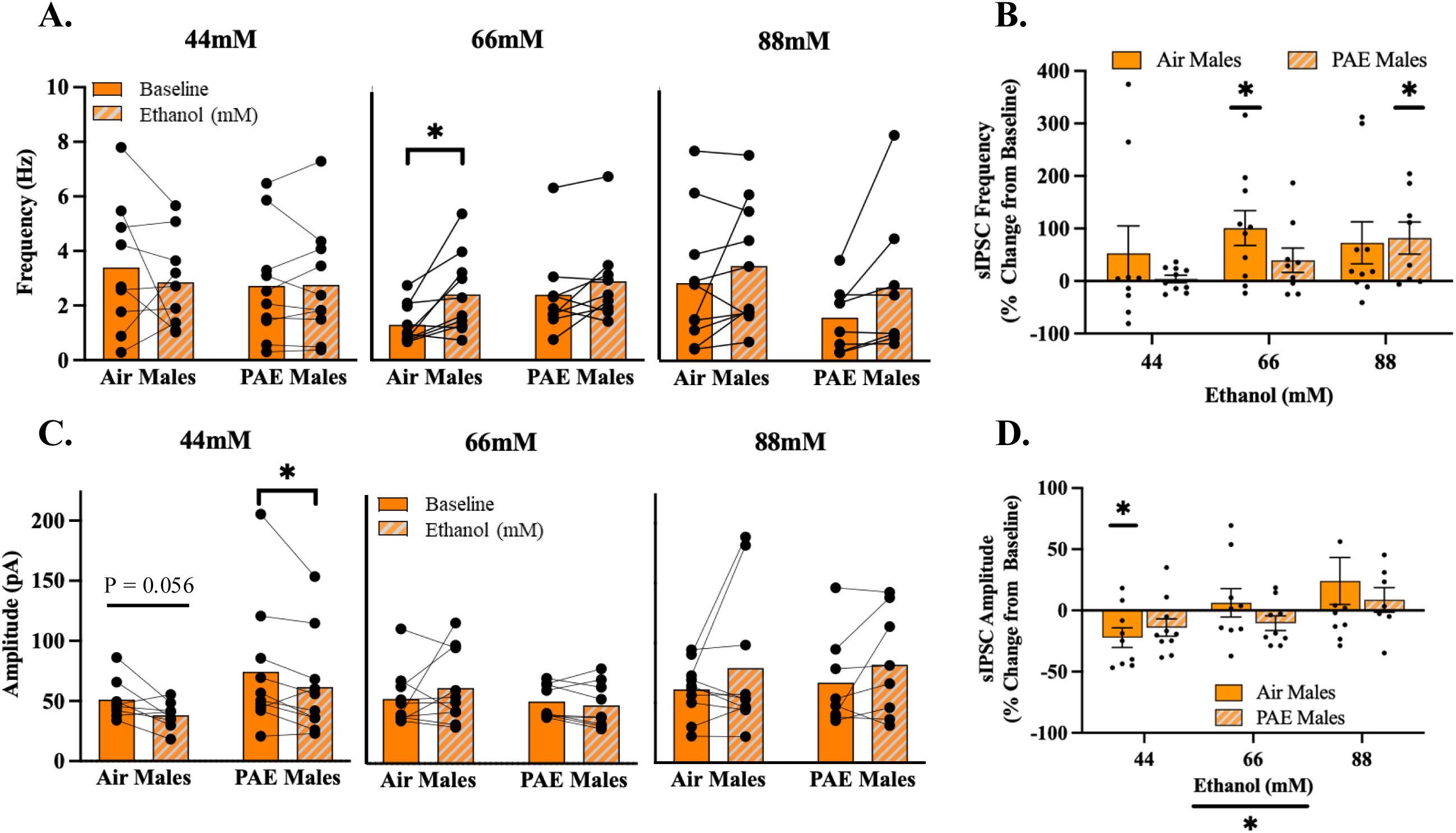
Acute EtOH action in the CeM – Males. (**A**) In air males, we observed an effect of acute alcohol on raw sIPSC frequency at 66 mM EtOH only (air: n = 8 cells from 8 animals; PAE: n = 9 cells from 9 animals), this was mirrored in % change from baseline with an increase of sIPSCs in air males at 66 mM EtOH (**B**). Though we did not see a significant change in the raw sIPSCs (**A**), sIPSC frequency was increased from baseline in PAE at 88 mM EtOH (n = 8 cells from 8 animals) (**B**). (**C**) There was a trend observed for a decrease in raw sIPSC amplitude in air males (n = 9 cells from 9 animals), and a significant decrease in raw sIPSC amplitude in PAE males (n = 10 cells from 10 animals). (**D**) 44 mM EtOH significantly decreased sIPSC amplitude from baseline in air males (n = 9 cells from 9 animals), (**D**) There was a main effect of concentration in the dose response curve. All data was tested for normal distribution and the appropriate parametric tests were used. * denotes p ≤ 0.05. Bars represent standard error of mean.

When examining raw changes in sIPSC amplitude, while we observed a strong trend of decreased sIPSC amplitude following application of 44 mM EtOH in air males (t (7) = 2.288, *p* = 0.056; **Figure 3C**), there were no effects at 66 mM (t (9) = 0.959, *p* = 0.363; **Figure 3C**) or 88 mM (t (9) = 1.228, *p* = 0.251; **Figure 3C**). Similar to air exposed males, 44 mM EtOH significantly decreased sIPSC amplitude in PAE males (t (9) = 2.325, *p* = 0.045; **Figure 3C**), with no significant effect of EtOH observed at 66 mM (t (8) = 1.096, *p* = 0.305; **Figure 3C**) or 88 mM (t (7) = 1.454, *p* = 0.189; **Figure 3C**). When analyzed as % change from baseline, we observed a significant decrease in sIPSC amplitude in air males at 44 mM EtOH (t (8) = 2.767, *p* = 0.024; **Figure 3D**), which was not present at 66 mM (t (8) = 0.543, *p* = 0.602; **Figure 3D**), or 88 mM (t (9) = 1.257, *p* = 0.241; **Figure 3D**). Conversely, there was no effect of EtOH on the sIPSC amplitude in PAE males at any concentrations (44 mM - t (9) = 1.974, *p* = 0.080; 66 mM - t (8) = 1.739, *p* = 0.120; 88 mM - t (6) = 0.861, *p* = 0.422; **Figure 3D**).

Finally, we assessed if mPAE shifted the dose response curve for males. A two-way ANOVA revealed no differences in the dose response curve when examining sIPSC frequency as a function of exposure (*F* (1, 50) = 1.475, *p* = 0.230) or concentration (*F* (2, 50) = 1.214, *p* = 0.306), and there was no interaction (*F* (2, 50) = 0.594, *p* = 0.556) (**Figure 3B**). Interestingly, a two-way ANOVA revealed a main effect of concentration on sIPSC amplitude (*F* (2, 48) = 4.297, *p* = 0.019). However, there were no differences as a function of exposure (*F* (1, 48) = 0.687, *p* = 0.411) nor an interaction (*F* (2, 48) = 0.738, *p* = 0.483) observed in sIPSC amplitude for the dose response curve (**Figure 3D**).

### 3.6 G12 mPAE alters acute alcohol modulation of GABA transmission in female CeM neurons

Similar to males, we examined the effect of PAE on the actions of acute alcohol in females. When examining raw changes in sIPSC frequency, we observed no changes in either air or PAE females at 44 mM (Air - W = 15.00, *p* = 0.492; PAE - t (8) = 1.732, *p* = 0.122), 66 mM (Air - t (9) = 0.705, *p* = 0.499; PAE - t (9) = 0.354, *p* = 0.732), or 88 mM (Air - t (9) = 0.734, *p* = 0.647; PAE - t (9) = 1.818, *p* = 0.103; **Figure 4A**). However, when sIPSC frequency was analyzed as % change from baseline, we observed a modest, but significant increase in PAE females at 88 mM EtOH (t (9) = 2.262, *p* = 0.050; **Figure 4B**), with a trend of increased sIPSCs at 44 mM EtOH (t (8) = 2.162, *p* = 0.063; **Figure 4B**), and no differences observed at 66 mM (t (8) = 0.954, *p* = 0.368; **Figure 4B**). In contrast, we saw no differences in % change from baseline sIPSCs frequency in air females at 44 mM (t (9) = 0.739, *p* = 0.479; **Figure 4B**), 66 mM (t (9) = 1.414, *p* = 0.191; **Figure 4B**), or 88 mM (t (9) = 1.427, *p* = 0.187; **Figure 4B**). Air females exhibited no raw changes in sIPSC amplitude in response to 44 mM (W = −9.000, *p* = 0.695; **Figure 4C**), 66 mM (t (9) = 0.571, *p* = 0.582; **Figure 4C**), or 88 mM EtOH (W = 3.000; *p* = 0.922; **Figure 4C**). No significant changes were observed in PAE females for sIPSC amplitude at 44 mM (t (7) = 1.047, *p* = 0.330; **Figure 4C**), 66 mM (t (9) = 0.280, *p* = 0.786; **Figure 4C**), or 88 mM (t (9) = 0.750, *p* = 0.473; **Figure 4C**). When analyzed as % change from baseline, there were no changes observed in sIPSC amplitude at any concentration in air (44 mM - t (9) = 0.263, *p* = 0.798; 66 mM - t (9) = 0.121, *p* = 0.906; 88 mM - t (9) = 0.614, *p* = 0.555; **Figure 4D**) or PAE females (44 mM - t (8) = 1.378, *p* = 0.205; 66 mM - t (9) = 0.147, *p* = 0.887; 88 mM - t (9) = 0.927, *p* = 0.378; **Figure 4D**).

**Figure 4.**
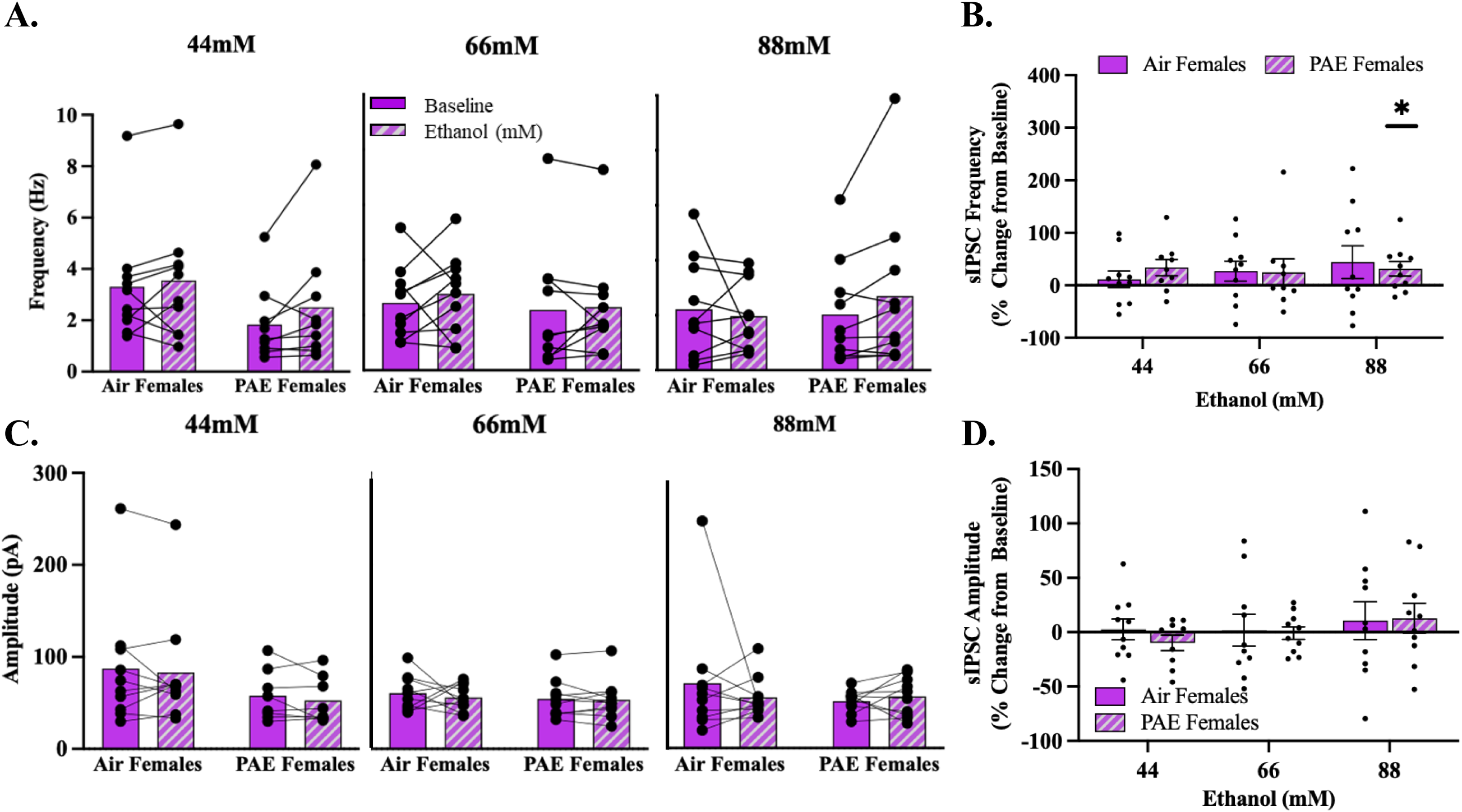
Acute EtOH action in the CeM – Females. (**A**) There were no differences in raw sIPSCs in air females or PAE females across concentrations. (**B**) All concentrations of EtOH did not alter sIPSC frequency in air females when measured as % change from baseline, but sIPSC frequency was increased in PAE females at the highest concentration (88 mM EtOH; n = 10 cells from 10 animals). (**C**) There were no differences in raw sIPSC amplitude across concentrations in either air or PAE females. (**D**) We observed no shifts in the dose response curve for sIPSC amplitude in either air or PAE females. All data was tested for normal distribution and the appropriate parametric tests were used. * denotes p ≤ 0.05. Bars represent standard error of mean.

Finally, we observed no shifts in the dose response curve in frequency (**Figure 4B**) or amplitude (**Figure 4D**) as a function of exposure (frequency - *F* (1, 52) = 0.018, *p* = 0.893; amplitude - *F* (1, 53) = 0.185, *p* = 0.669), concentration (frequency - *F* (2, 52) = 0.287, *p* = 0.752; amplitude - *F* (2, 53) = 0.846, *p* = 0.435), or an interaction (frequency - *F* (2, 52) = 0.358, *p* = 0.701; amplitude - *F* (2, 53) = 0.180, *p* = 0.836) between these effects.

## 4.0 Discussion/Conclusion

We aimed to assess neuronal and synaptic mechanisms within the CeM that may contribute to alterations in alcohol-related behaviors in a rat model of mPAE. We found, under basal conditions, that PAE males required more current to elicit an AP, and that PAE females showed reduced GABA transmission, indicated by reduced sIPSC frequency. Furthermore, we wanted to characterize the actions of acute EtOH in CeM neurons in adolescent animals, an area of research that has remained largely unexplored. We found that EtOH increased GABA release in control males at the tested middle concentration (66 mM EtOH), yet GABA release was increased in PAE males only at the highest concentration (88 mM EtOH). Furthermore, we saw postsynaptic effects at 44 mM EtOH in PAE males. Interestingly, 44 mM EtOH also decreased the amplitude in air males from baseline, though this did not reach significance. Consistent with previous reports in EtOH naïve adults, adolescent air females were insensitive to the effects of acute alcohol at all of the concentrations tested. However, as we hypothesized, we saw a significant increase in GABA release at the highest concentration (88 mM EtOH) in PAE females. Taken together, these data support our hypothesis that PAE produced sex-specific alterations in synaptic activity and neuronal function.

In our current study, we observed an increased rheobase, a measure of excitability which assesses the amount of current needed to elicit an action potential, in PAE males relative to control males. While this does suggest that these cells are hypoexcitable, we did not observe the same PAE induced neuronal alterations as previously found in our lab (Rouzer and Diaz, 2022) which may be due, in part, to differences in age of the animals tested. For example, we previously observed reduced excitability in PAE adolescent males (maximum of ~4-5 current-induced AP in control males and ~2-3 in PAE males) that likely ranged from peri-pubertal (P40-48). In the current study, we assessed excitability in late adolescents (P45-55), likely to be post-pubertal, where even control males only exhibited a maximum of ~2 current-induced AP. Interestingly, we previously found that adult males exhibit reduced excitability compared to adolescents, with a maximum of ~2 current-induced AP in adult males (Rouzer and Diaz, 2021), similar to what we have observed here in both PAE and air males. It may be that the neurophysiological maturation of these cells peaks in late adolescence before the age we previously considered ‘adulthood’, which may contribute to differences between our current and previous PAE studies. In addition, we previously reported a PAE-induced depolarization of CeM neurons (PAE males: ~ −69 mV; PAE females: ~ −67 mV)(Rouzer and Diaz, 2022), which was not observed here (PAE males: ~ −46 mV; PAE females ~ −43 mV). One likely possibility is the differences in internal solution used across studies prevented us from capturing PAE-induced differences that we previously reported. In previous studies, a K-gluconate-based internal solution was used to assess excitability of cells, however, we used a KCl-based internal solution in order to assess spontaneous activity and excitability within the same cell. Regarding synaptic activity, we found a reduction in basal GABA transmission in PAE females, indicated by reduced sIPSC frequency, yet no change in sIPSC amplitude, suggesting reduced presynaptic GABA release. Interestingly, we previously reported a reduction in sIPSC frequency in adolescent PAE males (Rouzer and Diaz, 2022). These differences between the current and past study may also be related to the difference in age range, which is something that should be further investigated. Regardless, given that the CeM is the major output of the CeA and contains mostly GABAergic interneurons and projection neurons, alterations in GABA transmission onto CeM neurons ultimately affects local and long-range inhibitory tone. Thus, the current finding showing a decrease in local inhibitory tone may cause increased GABA release downstream. Conversely, a reduction in long-range inhibitory tone could cause hyperexcitability of downstream targets. While we did not have the capabilities to identify the CeM neurons we recorded as either local or projection, either situation could contribute to PAE-induced alcohol misuse. Future studies should more directly examine PAE effects in projection-specific neurons to downstream targets relevant to alcohol-related behaviors.

The actions of acute EtOH have been well established in multiple brain regions of naïve male animals (Harrison et al., 2017). Concentrations as low as 10 mM ranging to 50 mM increase sIPSCs in the cerebellum, with similar effects shown in the hippocampus, whereas increases in GABA transmission were concentration dependent in the ventral tegmental area (Harrison et al., 2017). In the CeA, Roberto and colleagues have established that a moderate concentration of EtOH (44 mM) increases GABA transmission in adult naïve males (Kirson et al., 2021; Roberto et al., 2003; Roberto et al., 2004). To our knowledge this is the first study to examine how acute EtOH affects GABA transmission specifically in the CeM of late adolescent animals (P45-55) and how PAE may alter acute EtOH actions. Air males exhibited the largest increase in sIPSC frequency in response to 66 mM EtOH, suggesting increased GABA release, with no significant change in GABA transmission at 44 mM or 88 mM. It is worth noting that while 88 mM EtOH did generally increase sIPSC frequency in air males, despite having a sufficient sample size, the effect did not reach significance. Based on the spread of the individual data points, it is clear that this lack of significant effects was likely due to the variability. Interestingly, PAE males were sensitive to acute EtOH effects only at the highest concentration, with 88 mM increasing sIPSC frequency. While this is certainly suggestive of a shift in concentration response, analysis of the concentration response curve did not reveal a significant main effect of exposure. Although we did not explore mechanisms contributing to these PAE-induced differences in sensitivity to acute ethanol, it has been reported that EtOH induced increase of GABA release is mediated, in part, by the corticotropin releasing factor receptor type I (CRFR1) in the CeM of adult males (Nie et al., 2004; Nie et al., 2009; Roberto et al., 2010). Our lab previously found a PAE-induced reduction of CRFR1+ cells (Rouzer and Diaz, 2022), which may contribute to PAE males requiring higher concentrations of EtOH to increase GABA release. Studies are currently underway to test the impact of PAE on interactions between acute ethanol and CRFR1 mechanisms. In addition to acute EtOH actions on presynaptic mechanisms, 44 mM produced a significant change in sIPSC amplitude in PAE and air males, suggestive of potential postsynaptic actions of ethanol. It is possible that this concentration of ethanol may acutely alter receptor kinetics or sensitivity independent of PAE exposure, an effect that has been previously shown in adult naïve males (Roberto et al., 2003).

With females, our findings align with reports made by Kirson *et al*. (2021), in that females are insensitive to acute EtOH action at any concentration (44, 66, 88 mM) in late adolescence. Interestingly, we observed a significant increase in sIPSC frequency in PAE females at the highest tested ethanol concentration (88 mM), suggesting that mPAE may produce alterations similar to chronic EtOH exposure in adults (Kirson et al., 2021). This could be due, in part, to sex differences in corticotropin-releasing factor (CRF) signaling. A previous study showed that CRF potentiated GABA in the same manner as acute EtOH through its activity at the CRFR1 in adult naïve males (Rodriguez et al., 2022). However, while adult naïve female rats are unresponsive to the actions of CRF at multiple concentrations (100, 200, 400 nM), the highest concentration of CRF increased GABA transmission in alcohol-dependent females (Rodriguez et al., 2022). Interestingly, we have shown an upregulation in CRFR1+ neurons in PAE adolescent females (Rouzer and Diaz, 2022) similar to the increase in CRF-mediated effect observed in alcohol dependent adult females (Rodriguez et al., 2022). Taken together, our data in combination with other studies suggest that the neurophysiological maturation of these cells may peak in late adolescence (P45-55) before the age that we have considered ‘adulthood’, which likely contributes to the lack of responsivity to acute EtOH. However, whether acute ethanol alters GABA transmission in the CeM of younger females has yet to be tested. Additionally, our data suggests that mPAE may produce a heightened sensitivity in the CRF system of adolescent females similar to what is observed following chronic alcohol exposure in adult females. Further investigation into this system is needed to understand the mechanisms contributing to sex differences in acute ethanol and CRF actions.

PAE is a global insult, therefore, our G12 PAE may alter many different systems that are developing at the time of exposure. Our study revealed significant neurophysiological differences within the CeM between PAE and control animals that may contribute to the increased risk of alcohol misuse and dependence. However, we acknowledge that the CeM is not the only structure involved and it may be that these neurobiological mechanisms do not completely explain the heightened risk of developing AUD in individuals with FASD. In conclusion, it is imperative that we continue to inform the public about the risk of alcohol consumption during pregnancy. A single G12 mPAE produced several neurophysiological changes in late adolescent animals, including a change in the concentrations of ethanol that acutely modulate GABA transmission within the CeM in both males and females. Adolescence is a vulnerable developmental period when individuals engage in risk taking behaviors, such as misuse of alcohol (Spear, 2000). Therefore, it is essential to understand how PAE induced neurophysiological changes contribute to alcohol-related behaviors as well as how alcohol’s action in the brain is altered by PAE. Thus, continued research in this field has the potential to unveil underlying mechanisms involved in PAE induced substance misuse.

